# Interfacing Seurat with the R tidy universe

**DOI:** 10.1101/2021.03.26.437294

**Authors:** Stefano Mangiola, Maria A Doyle, Anthony T Papenfuss

## Abstract

**Motivation:** Seurat is one of the most popular software suites for the analysis of single-cell RNA sequencing data. Considering the popularity of the tidyverse ecosystem, which offers a large set of data display, query, manipulation, integration and visualisation utilities, a great opportunity exists to interface the Seurat object with the tidyverse. This gives the large data science community of tidyverse users the possibility to operate with a familiar grammar.

**Results:** In order to provide Seurat with a tidyverse-oriented interface without compromising on efficiency, we developed tidyseurat, a light-weight adapter to the tidyverse. Cell information is automatically displayed as a tibble abstraction, which interfaces Seurat with dplyr, tidyr, ggplot2 and plotly packages powering efficient data manipulation, integration and visualisation. Iterative analyses on data subsets is enabled by interfacing with the popular nest-map framework.

**Availability and implementation:** The software is freely available at cran.r-project.org/web/packages/tidyseurat/ and github.com/stemangiola/tidyseurat

**Contact:** Stefano Mangiola and Anthony T Papenfuss.

## Introduction

Nucleotide sequencing at the single-cell resolution level has proven to be a disruptive technology that is revealing unprecedented insights into the role of heterogeneity and tissue microenvironment in disease (Xiao *et al.,* 2019; Keil *et al.*, 2018). Single-cell RNA sequencing data allows the robust characterisation of tissue composition (Abdelaal *et al.*, 2019), the identification of cellular developmental trajectories (Gojo *et al.*, 2020; Chen *et al.*, 2019; Van den Berge *et al.*, 2020; Saelens *et al.*, 2019), and the characterisation of cellular interaction patterns (Cabello-Aguilar *et al.*; Kumar *et al.*, 2018; Shao *et al.*, 2020). In recent years, the scientific community has produced a large number of computational tools for the analysis of such data (Butler *et al.*, 2018; Lun *et al.*, 2016; McCarthy *et al.*, 2017). One of the most popular of these, Seurat (Butler *et al.*, 2018; Stuart *et al.*, 2019), stores raw and processed data in a highly optimised, hierarchical structure (Figure 1A). This structure is displayed to the user as a summary of its content. The user can extract and interact with the information contained in such a structure with Seurat custom functions.

**Figure 1.**
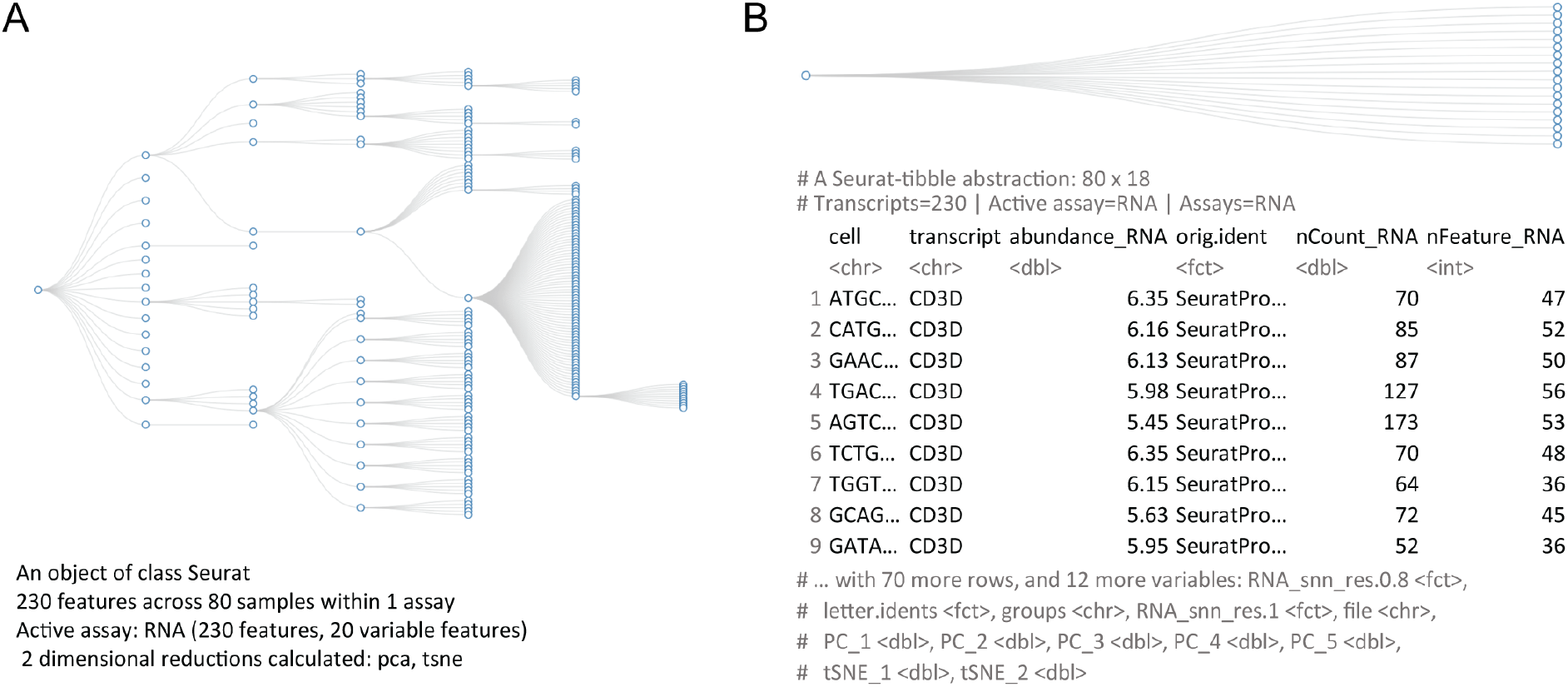
Comparison between the data structure (Cui, 2020) (top; abstracted tibble for tidyseurat) and the information presented to the user (bottom) for Seurat (panel A) and tidyseurat (panel B; including transcript information). The data set underlying these visualisations is a subset of a peripheral blood mononuclear cell fraction provided by 10X (10xgenomics.com/).

Machines and humans often have orthogonal needs when interacting with data. While machines prioritise memory and computation efficiency and favours data compression, humans prioritise low-dimensional data display, and direct and intuitive data manipulation. Considering that low-dimensionality data representation often requires redundancy, it is challenging to balance all priorities in a unique data container. Separating roles between the back-end data container and the front-end data representation is an elegant solution for ensuring both transparency and efficiency. The scientific community has tackled this issue by offering visual and interactive representation of Seurat single-cell data containers. For example, Cerebro (Hillje *et al.*, 2020) is a Shiny-based standalone desktop application (Web Application Framework for R [R package shiny version 1.5.0], 2020) that enables the investigation and the inspection of pre-processed single-cell transcriptomics data without requiring bioinformatics experience. This application can import and export Seurat data containers. Similarly, BioTuring (BioTuring INC) offers a web visual interface for facilitating data analysis by scientists without coding experience. NASQAR (Yousif *et al.*, 2020) (GitHub.com/nasqar) enables interactive analysis of a wide variety of genetic data including single-cell RNA sequencing data from Seurat. Single Cell Viewer (SCV) (Wang *et al.*) is an R shiny application that offers users rich visualization and exploratory data analysis options for single cell datasets, including Seurat. Although these tools allow an intuitive data representation and analysis, they are not fully programmable and therefore not suitable for reproducible research. Moreover, they are hardly expandable and cannot easily incorporate all tools that the scientific community publishes in R data analysis repositories such as CRAN (Ripley, 2001) and Bioconductor (Huber *et al.*, 2015).

Recently, efforts have been made toward the representation and manipulation of data using the concept of data tidiness (Wickham *et al.*, 2019). This paradigm allows the organisation of information as a two-dimensional, highly flexible table (referred to as tibble, a type of a data frame), with variables oriented in columns and observations oriented in rows. This new standard has become extremely popular in many fields of data science. The application of tidiness principles would be extremely powerful applied to single-cell transcriptomic data; because it directly captures how biological data measurements relate to experimental design and metadata (e.g. technical and clinical properties of transcripts, cells and biological replicates). The shift from a compressed and hierarchical to a tabular data representation of cell-(by default) and/or transcript-related information has the immediate advantages for scientific awareness, and enables the direct interfacing with a large ecosystem of tidy-oriented APIs for data manipulation and visualisation, thus facilitating data analysis and reproducibility for researchers across a wide spectrum of computational literacy. For example, tidyseurat allows to display, plot, modify, join or delete information, filter, subsample, nest and map functions, summarise information of a Seurat object without leaving the tidyverse syntax and without the need of package specific syntax. This is particularly powerful in perspective of a tidy counterpart for SingleCellExperiment (Amezquita *et al.*, 2020) objects, moving toward a unified interface for single-cell data containers. As for comparison, although the indirect interface between Seurat objects and the tidyverse is possible, it requires intermediate steps in order to extract information that can be passed to downstream APIs. For example, building a custom plot that integrates reduced dimensions with cell-wise annotations (e.g. library size, mitochondrial transcription, cell-type identity) requires first an integration of multiple data frames (e.g. included in the slots metadata, and reductions) with custom routines (e.g. direct querying for metadata and Embeddings for reduced dimensions).

Here we present tidyseurat, an adapter that interfaces Seurat, the most popular single-cell RNA sequencing data analysis tool, with tidyverse, the most popular R data analysis framework. Although the underlying data container is Seurat’s, when displayed on screen the cell-wise information (normally hidden from the user the the object is displayed on screen) is organised as a tibble abstraction (Figure 1B). Tidyseurat includes adapters to the vast majority of methods included in dplyr (Hadley Wickham *et al.*, 2019), a powerful grammar of data manipulation; tidyr (Mailund, 2019) a large collection of methods for data reshaping and grouping; ggplot2 (Wickham *et al.*, 2016), the most popular R visualisation tool; and plotly (Inc, 2015), a powerful tool for interactive visualisations. As a result, the user can perform efficient analyses using Seurat (and Seurat-compatible software), while visualising, manipulating, integrating and grouping the data using tidyverse (-compatible for plotly) software. This package is aimed at analysts of single-cell data who favors the use of tidyverse and Seurat. Tidyseurat is part of a larger ecosystem called tidytranscriptomics that aims to bridge the transcriptomics and tidy universes (github.com/stemangiola/tidytranscriptomics).

### System and methods

#### Data user interface

Tidyseurat abstracts the complexity of the data container and provides a friendlier interface for the user. This abstraction can be obtained by applying the method ‘tidy()’ to a Seurat object. Tidyseurat implements an improved data display method (replacing the Seurat ‘show’ method) mapping the cell-wise information into a user-friendly table. By default, cell-wise information is displayed to the user (e.g. cell-cycle phase, cluster and cell-type annotation), leaving the transcript information available upon request using the ‘join_transcripts’ function. This function adds transcript identifiers, transcript abundance and transcript-wise information (if present; e.g. gene length, genomic coordinates and/or functional annotation) as additional columns. Cell-wise information is prioritised over transcript-wise information on the rationale that it is more often directly queried.

The tidyseurat tibble abstraction includes two types of columns, columns that can be interacted with and modified, and columns that are view only. The editable columns are part of the cell metadata, while the view-only columns are those that are calculated such as reduced dimensions (e.g. principal component and UMAP dimensions). The default integration of all cell-wise information in one tibble representation, including reduced dimensions facilitated data visualisation, filtering and manipulation. To allow the manipulation and plotting of the data using the tidyverse ecosystem, the dplyr, tidyr, ggplot2, and the tidyerse-compatible plotly routines have been adapted to work seamlessly with the back-end Seurat data structure, allowing the user to operate as if it was a standard tibble. This abstraction strategy allows the data to appear as a tibble for end-users and the tidyverse (Table 1) but appear as a Seurat container for any other algorithm, thus preserving full backward compatibility (Figure 2).

**Table 1.** Example of a tidyseurat table. Pre-existing cell-wise annotation and new calculated information is all coexisting in a unique table.

| # A Seurat-tibble abstraction: 8033 x 11 |  |  |  |  |  |  |  |  |  |  |
| --- | --- | --- | --- | --- | --- | --- | --- | --- | --- | --- |
| # Transcripts=1000 \| Active assay=SCT \| Assays=RNA, SCT |  |  |  |  |  |  |  |  |  |  |
| Cell | Total count | Total transcripts | PC1 | PC2 | PC... | UMAP2 | UMAP2 | UMAP... | Cluster | Cell type |
| cell_1 | 10456 | 450 | -1.23 | -2.11 | ... | -3.47 | 3.51 | ... | 1 | T cell |
| cell_2 | 2088 | 400 | 0.98 | 3.09 | ... | -1.59 | -2.8 | ... | 2 | B cell |
| cell_3 | 11309 | 699 | 5.55 | -0.02 | ... | 1.26 | 0.39 | ... | 5 | Monocyte |
| cell_4 | 8791 | 423 | -5.42 | 1.12 | ... | -4.42 | -3.63 | ... | 1 | Monocyte |

**Figure 2.**
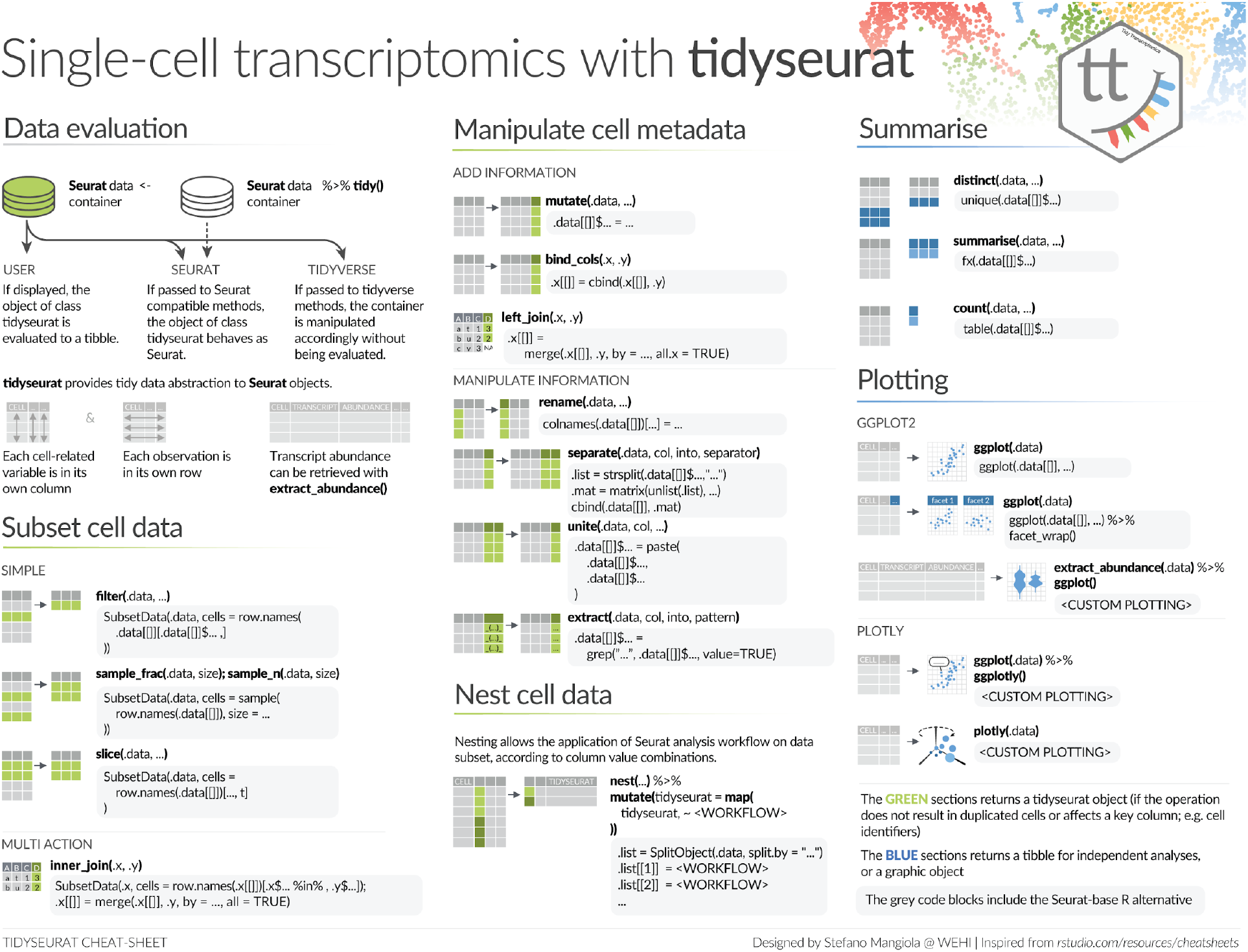
Cheatsheet of the tidyverse functionalities that tidyseurat enables for Seurat objects. This cheatsheet provides examples of the alternative tidyverse and Seurat syntax. The green colour scheme includes procedures that output a tidyseurat, if: (i) do not lead cell duplication; and (ii) key columns (e.g. cell identifier) are not excluded, modified nor renamed (e.g., through a select, mutate and/or rename commands). In this case, a table (rather than an abstraction) is returned for independent analysis and visualisation. The blue colour scheme includes procedures that return tibble tables for independent analyses and plotting. The grey-shaded boxes include the alternative code utilising Seurat and base-R.

#### API user interface

Thanks to R’s S4 class inheritance, tidyseurat objects can be operated by any Seurat compatible algorithms. The seamless integration with the tidyverse is obtained through adapters for the vast majority of methods in the packages dplyr, tidyr, ggplot2; as well as plotly (Figure 2). These methods can be separated into three groups based on the action that they perform on the back-end Seurat container. Methods such as ‘mutate’, ‘left_join’, ‘separate’, ‘unite’ and ‘extract ‘, ‘select’ manipulate or subset the information present in the cell-wise metadata. Methods such as ‘slice’, ‘filter’, ‘sample_n’, ‘sample_frac’, ‘inner_join’ and ‘right_join’ subset specific cells based on a wide range of criteria. Methods such as ‘bind_rows’ join two or more datasets. All these methods return tidyseurat tables if those procedures do not lead cell duplication and if key columns (e.g. cell identifier) are not excluded; otherwise a tibble is returned for independent analyses. Another group of functions such as ‘summarise’, ‘count’, ‘distinct’, ‘join_transcripts’ and pull return a tibble and an array respectively by design, for independent analyses. Tidyverse-compatible visualisation methods include ggplot and plotly. These methods operate on the tibble abstraction of the data. Such abstraction allows the application of the nest-map tidyverse framework. Briefly, nesting enables tables to be divided into subsets according to any combination of columns and to nest them into a column; the map function allows to iteratively apply operations across subsets. Applied to single-cell data containers, this tool confers great robustness, flexibility and efficiency.

#### Algorithm and implementation

To demonstrate the use of tidyseurat, we provide as example an integrated analysis of peripheral blood mononuclear cells from public sources. We show the main steps of a common workflow, along with code examples (Figure 3) and tidyverse-compatible visualisations (Figure 4). As an example, we show how data manipulation and filtering can result in a three folds reduction in coding lines and a decrease in temporary variables compared to Seurat alone (Supplementary code chunk 1).

**Figure 3.**
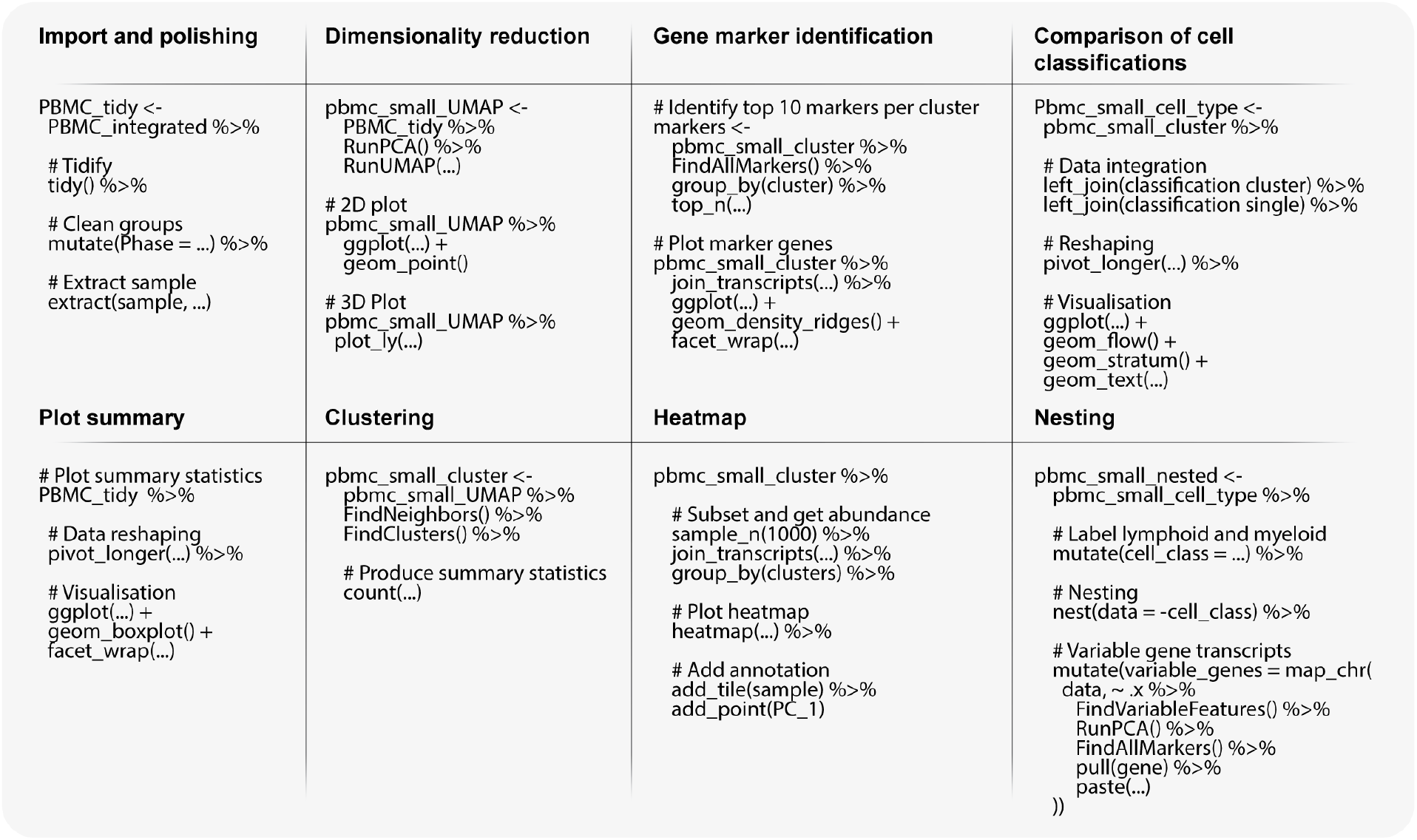
Pseudo-code representing the procedures for the analysis of single-cell RNA sequencing data analysis using integrating Seurat and tidyverse functions through tidyseurat.

**Figure 4.**
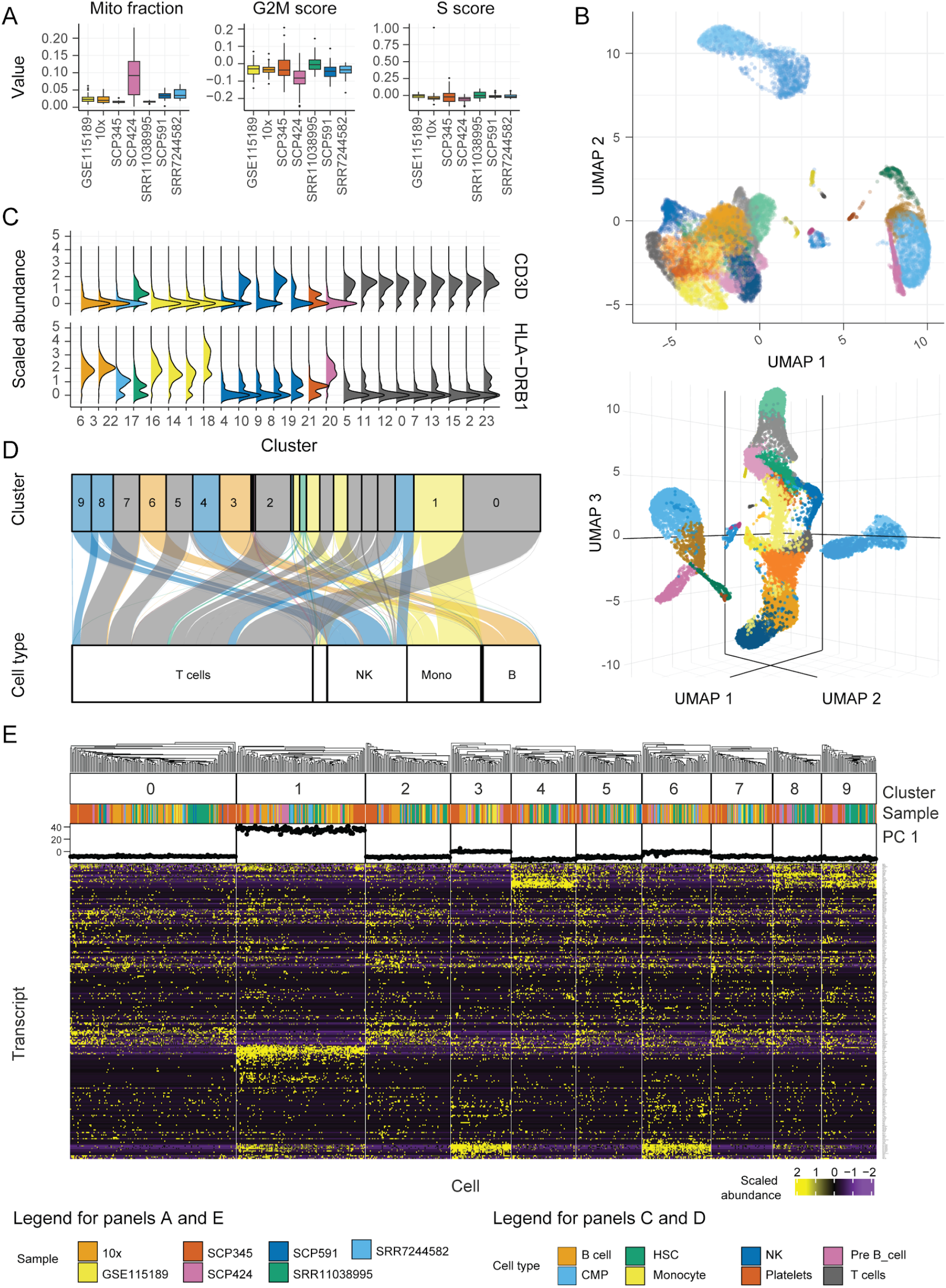
Tidyverse-compatible libraries offer powerful, flexible and extensible tools to visualise single-cell RNA sequencing data. Natively interfacing with such tools expands the possibilities of the user to learn from the data. Graphical results of the example workflow, integrating Seurat and tidyverse with tidyseurat. **A.** Sample-wise distribution of biological indicators; including proportion of mitochondrial transcripts and cell-cycle phase scores. For optimum visualisation, a 20% subsampling was performed on the cell set. **B.** Cells mapped in two- and three-dimensional UMAP space. The default integration of reduced dimensions, together with other cell-wise information, in a tibble abstraction facilitates such visualisation. **C.** Distribution of transcript abundance for some marker genes, identified for each cluster identified by unsupervised estimation. Cells mapped in two-dimensional Uniform Manifold Approximation and Projection (UMAP) space. **D.** Mapping of cells between the cell- or cluster-wise methods for cell-type classification. Only large clusters are labelled here. The colour scheme refers to cell types classified according to clusters. The bottom containers refer to the classification based on single cells. **E.** Heatmap of the marker genes for cell clusters, produced with tidyHeatmap, and annotated with data source and the first principal component. Only the ten largest clusters are displayed. The integrated visualisation of transcript abundance, cell annotation, and reduced dimensions is facilitated by the ‘join_transcripts’ functionality and by the default complete integration of cell-wise information (including reduced dimensions) in the tibble abstraction.

#### Data import, polishing and exploration

The single-cell RNA sequencing data used in this study consists of seven datasets of peripheral blood mononuclear cells, including GSE115189 (Freytag *et al.*, 2018), SRR11038995 (Cai *et al.,* 2020), SCP345 (singlecell.broadinstitute.org), SCP424 (Ding *et al.*, 2019), SCP591 (Karagiannis *et al.*, 2020) and 10x-derived 6K and 8K datasets (support.10xgenomics.com/). In total, they include 50706 cells. Data exploration is a key phase of any analysis workflow. It includes data visualisation and production of summary statistics, in combination with dimensionality reduction and data scaling. The Seurat data container is abstracted to its tidy representation with the command ‘tidy’, and the polishing of cell-wise annotations (cell-cycle phase and sample name) can be achieved with tidyverse commands (Figure 3, Import and polishing; Supplementary code chunk 2). Key cell properties included in the resulting table (e.g. proportion of mitochondrial transcripts and cell-cycle phase; Figure 4A) can be visualised in a faceted and integrated fashion using common tidyverse tools (Figure 3, Plot summary). This visualisation facilitates quality control, helping identify potential low-quality samples such as SCP424 (Figure 4A).

#### Dimensionality reduction

Dimensionality reduction is a key step of single-cell RNA sequencing data analysis, as it allows visualising cell heterogeneity in a plot (Figure 3, Dimensionality reduction). Beside principal component analysis (PCA) (Venables and Ripley, 2002), methods such as uniform manifold approximation and projection (UMAP) (McInnes *et al.*, 2018) are commonly used to better define local similarities, while preserving global distances. Seurat and tidyverse methods can be seamlessly integrated through tidyseurat for the calculation and visualisation of UMAP dimensions (Figure 3, Dimensionality reduction). The reduced dimensions are displayed as (view only) additional columns of the tidyseurat table. The use of tidyverse (Wickham *et al.*, 2019) for visualisation allows great customisation of two-dimensional plots (Figure 4B). The advantage of tidyseurat here is the presence of cell annotation and reduced dimension in the same data frame, that can be used for arbitrarily complex annotated visualisations (Supplementary code chunk 3). Three-dimensional plots can be produced effortlessly applying plotly (Sievert, 2020) on the tidyseurat tibble abstraction (Figure 4B). Visualising a third reduced dimension confers better awareness of cell heterogeneity and clustering. Dimensionality reduction shows 3 main cell clusters and a minor intermediate cell cluster (Figure 4B, top). The main cluster (bottom-left) includes 69% of all cells (Mangiola, 2020). The display of the third UMAP dimension in an interactive environment gives an additional perspective on cell heterogeneity and compared to only calculating and visualising the first two (Figure 4B, bottom).

#### Clustering and marker genes identification

Unsupervised clustering based on cell transcription is essential to quantitatively define self-similar groups of cells. Similarly to previous procedures, Seurat and tidyverse commands can be concatenated though inference and visualisation steps (Figure 3, Clustering; Supplementary code chunk 4). The newly calculated cluster identities will be displayed as additional columns in the tidyseurat table. The cell clustering information can be used to identify the marker genes that are preferentially transcribed in each cell group (Figure 3, Gene marker identification). This information is key in order to define cell identities. Gene marker identification can be performed with Seurat, and transcript abundance distribution can be visualised for selected marker genes in a faceted and integrated manner using tidyverse (Figure 4C). The advantage of tidyseurat here is the ease of the integration of the transcript and cell information in the same data frame (through ‘join_transcripts’) for joint manipulation, filtering and visualisation (Supplementary code chunk 5). Cell-wise transcript abundance for marker genes can be also efficiently visualised using a heatmap. While it is possible to use the Seurat integrated heatmap function (DoHeatmap), the tidyverse-style heatmap method (Mangiola and Papenfuss, 2020), tidyHeatmap, allows for more flexibility. For example, several cell-wise data (e.g. principal components) can be added as annotations, choosing among several visualisations (e.g. tile, point, line and bar; Figure 4E). The integration of diverse information facilitates quality check and curation. Shared-nearest-neighbour (SNN) method (Ertöz *et al.*, 2003) for unsupervised clustering identified 24 cell clusters with default settings. The biggest cluster includes the 17% of cells. The biggest supercluster including 69% of all cells encompasses 18 clusters.

#### Cell type inference

While the classification of cell clusters in cell-type categories can be performed manually through the analysis of marker genes, the automatic cell and/or cluster classification can represent a key first step in the process. Several methods are publicly available (Alquicira-Hernandez *et al.*, 2019; Nagendran *et al.*, 2018; Jaitin *et al.*, 2014; Tan and Cahan, 2019; Kim *et al.*, 2019). SingleR (Aran *et al.*, 2019) is a popular tool, able to classify both clusters and single cells using transcriptional references. While using cluster identity to drive the cell-type classification can benefit from data aggregation and improve the overall robustness of the inference, it relies on the goodness of clustering and on the assumption that cells within the same cluster are of the same type. On the contrary, single-cell classification avoids biases due to clustering but introduces challenges relative to the absence of data hierarchy. Using tidyseurat, the consistency between these two methods can be visually and quantitatively checked (Figure 3, Comparison of cell classification). The tidyverse-style alluvial visualisation is ideal to communicate the differences in classification with or without cluster information, and integrates with the tidy data structure (Bojanowski and Edwards, 2016; Kennedy and Sankey, 1898; Brunson, 2020) (Figure 4D). Using the Human Protein Atlas reference (Uhlén *et al.*, 2015), eight cell types were identified in total (including platelets, T-, B-, pre-B-, natural killer, monocyte, myeloid progenitor and hematopoietic stem cells). For both classification approaches (cluster- or cell-wise) the most abundant cell type was T-cells, including on average 51% of all cells. In total, the 9.4% of cells were classified differently between the two methods (Figure 4D).

#### Nesting

Subsetting the data according to sample, cell identity and/or batch is a common step of a standard analysis workflow. For example, grouping according to major cell subtypes (e.g. lymphocytes, myeloid and stromal cells) might be needed for independent analysis with improved resolution; performing independent analyses across biological replicates can be useful to assure that data integration is not creating artifacts; or similarly, balanced subsampling across biological replicates might be needed for an unbiased visualisation of reduced dimensions. This can be obtained by manually splitting the data into subsets according to a variable and iteratively applying procedures to each subset. Tidyverse gives a more powerful and intuitive framework to perform such operations on tibbles. The functions nest (Mailund, 2019) allows to nest data subsets into a table column according to any combination of variables, and map (Henry and Wickham, 2018) allows iterating procedures across such subsets without leaving the clear and explicit tibble format. An example is shown (Table 2; Figure 3, Nesting) where (i) cell types are grouped in lymphoid and myeloid, and (ii) variable gene transcripts are independently identified for each of the two cell populations with an increased resolution, without the need to create any temporary variable.

**Table 2.** Example of a nested tidyseurat table, with gene markers calculated internally for each major immune cell type. This is obtained with the nest-map combination from tidyverse.

| # A tibble 2 x 3 |  |  |
| --- | --- | --- |
| Cell class | Data | Top Markers |
| lymphoid | <tidyseurat> | RPL34, RPS27, RPL32, RPS3A, RPL21, RPL31 |
| myeloid | <tidyseurat> | S100A8, S100A9, S100A12, VCAN, CYP1B1, CD14 |

## Discussion

Seurat is the most popular single-cell RNA sequencing data analysis workflow. It includes user-friendly methods for data analysis and visualisation. However, data query, manipulation and visualisation requires Seurat-specific functions. This ultimately limits critical data evaluation, scientific awareness and discoveries. The R data-science community has settled on a robust, consistent and modular data representation, which is referred to as tidy. Tidyseurat exposes the data contained in the complex hierarchical structure of a Seurat object in the form of a tidy table. As a result, the data is readily visible and transparent to the user, who can leverage the large computational and visualisation tidy ecosystem. Considering that tidyverse syntax and vocabulary (e.g. dplyr and tidyr) is becoming common knowledge, the domain-specific bioinformatic knowledge required to operate with Surat object is in practice greatly diminished. Most importantly, the full compatibility with the Seurat ecosystem is not compromised. By default, Seurat provides a wide range of custom methods for data plotting, where the data is internally extracted from its hierarchical container. The customizability of these methods is necessarily limited and achieved through setting command parameters. The tidyverse includes an increasing number of connected modules for data visualisation, that the transparent tidy data representation can leverage, eliminating the need of custom methods. The amount of information and heterogeneity within single-cell RNA sequencing data often requires data subsetting and reanalysis. For example, highly diverse broad cell populations such as lymphoid and myeloid are often subset and analysed independently to decrease the inference complexity and increasing resolution. While it is commonly required to manually subset the Seurat object, perform iterative analysis for each subset, and reintegrate the objects (if necessarily), the tidy abstraction enables the use of the nest-map paradigm. This elegant and powerful paradigm allows self-contained and robust iterative analysis on data subsets. Tidyseurat is a standalone adapter that improves analysis reproducibility and scientific awareness, in a user-friendly way, without changing the user’s familiar Seurat analysis workflow. As the display and manipulation is centered on cell-wise information by default, the use of tidyseurat does not add any perceptible overhead. This approach is particularly powerful in moving toward a unified interface for single-cell data containers together with a tidy counterpart for SingleCellExperiment objects. We anticipate that this data abstraction will be also the pillar of more extensive analysis-infrastructures based on the tidy paradigm, such as has happened for bulk RNA sequencing data (Mangiola *et al.*, 2021). In summary, tidyseurat offers three main advantages: (i) it allows tidyverse users to operate on Seurat objects with a familiar grammar and paradigm; (ii) it streamlines the coding, resulting in a smaller number of lines and fewer temporary variables compared with the use of Seurat only; and (iii) it provides a consistent user interfaces shared among other tidy-oriented tools for single-cell and bulk transcriptomics analyses (e.g. tidySingleCellExperiment and tidySummarizedExperiment). The package tidyseurat offers extensive documentation through methods description, vignettes (accessible typing browse Vignettes(“tidyseurat”)), and through workshop material (e.g. rpharma2020_tidytranscriptomics, ABACBS2020_tidytranscriptomics at github/stemangiola).

## Code availability

Tidyseurat is available on GitHub github.com/stemangiola/tidyseurat, and on CRAN cran.r-project.org/package=tidyseurat. The web page of the tidyseurat package is stemangiola.github.io/tidyseurat. The example code included in this manuscript is available as a markdown file at github.com/stemangiola/tidyseurat/vignettes. Seurat version 3 was used in this study.

## Author’s contribution

SM conceived and designed the method under the supervision of ATP. MD contributed to the comparative benchmark against base R standards. MD contributed with the software engineering and documentation. All authors contributed with manuscript writing.

## Acknowledgements

We thank the entire Bioinformatics Division of the Walter and Eliza Hall Institute of Medical Research for support and feedback.

